# Non-alcoholic steatohepatitis disrupts diurnal liver transcriptome rhythms in mice

**DOI:** 10.1101/2023.01.24.525228

**Authors:** Leonardo Vinicius Monteiro de Assis, Münevver Demir, Henrik Oster

## Abstract

**Background & Aims:** The liver ensures organismal homeostasis through modulation of physiological functions over the course of the day. How liver diseases such as non-alcoholic steatohepatitis (NASH) affects daily transcriptome rhythms in the liver remains elusive. To start closing this gap, we evaluated the impact of NASH on the diurnal regulation of the liver transcriptome in mice. Along this, we investigated how stringent consideration of circadian rhythmicity affects the outcomes of NASH transcriptome analyses.

**Approach & Results:** Comparative rhythm analysis of the liver transcriptome from diet-induced NASH and control mice revealed an almost 3h phase advance in global gene expression rhythms. Rhythmically expressed genes associated with DNA repair and cell cycle regulation showed increased overall expression and circadian amplitude. In contrast, lipid and glucose metabolism associated genes showed loss of circadian amplitude, reduced overall expression, and phase advances in NASH livers. Comparison of NASH-induced liver transcriptome responses between published studies revealed little overlap (12%) in differentially expressed genes (DEGs). However, by controlling for sampling time and using circadian analytical tools, a 7-fold increase in DEG detection was achieved compared to methods without time control.

**Conclusions:** NASH had a strong effect on circadian liver transcriptome rhythms with phase- and amplitude-specific effects for key metabolic and cell repair pathways, respectively. Accounting for circadian rhythms in NASH transcriptome studies markedly improves DEGs detection and enhances reproducibility.

## INTRODUCTION

Living beings are subject to rhythmic changes in environmental factors such as light or temperature. Mammals use internal circadian clocks to anticipate such regular changes and adjust physiological processes to meet environmental demands (1,2). Most biological processes show rhythmic activity such as heart rate, DNA repair, energy metabolism, and immunity (1,3–5). At the cellular level, circadian timekeeping is based on interlocked transcriptional-translational feedback loops (TTFLs) of clock genes and proteins. The clock TTFL orchestrates physiological rhythms through transcriptional programs of clock-controlled genes (3).

Circadian clocks are present in most cells and tissues including liver. Although the importance of circadian rhythms in liver physiology is well known, many basic and clinical studies often pay little attention to this factor when taking and analysing samples. A recent study evaluated the top-50 cited papers in 10 different fields between 2015 and 2019 showing that only 6.1% of the studies included time-of-day information (6). In recent years, the advances in omics techniques and analysis have clearly demonstrated that a substantial portion of the transcriptome, proteome, and metabolome shows rhythmic patterns (7–11). Therefore, a lack of temporal control may severely affect study outcome and reproducibility.

The prevalence of non-alcoholic fatty liver disease (NAFLD) is estimated at 25-30 % worldwide (12). NAFLD may progress to liver inflammation with tissue damage, a phenotype known as non-alcoholic steatohepatitis (NASH). If not treated, NASH will lead to cirrhosis or hepatocellular carcinoma (HCC). Unfortunately, no specific treatment for NAFLD is yet approved (13,14). NASH results from an imbalance of lipid metabolic processes like lipogenesis, beta-oxidation, and secretion of triglycerides from the liver, as well as endoplasmic reticulum stress, inflammation and insulin resistance, most of which are under circadian control (3,13,15–18). Disruption of circadian rhythms has been shown to hasten NASH and HCC in mice (19).

Several studies have identified gene signatures of NASH in mice and humans to identify prognostic markers for the different stages of NASH (20,21). Considering that a large portion of the liver transcriptome is rhythmic, we hypothesized that diurnal profiling of gene expression would critically improve detection of differentially expressed genes (DEGs) in this context.

## MATERIAL AND METHODS

### Mouse model and experimental conditions

Two to three months-old male and female wild type (C57BL/6J) mice were group-housed under 12h/12h light/dark conditions (LD, 200 – 400 lux) at 22 ± 2°C with food and water provided *ad libitum*. After 1 week of acclimatization under normal chow (NC, 5 % fat, 1314, Altromin, Germany) NASH was induced by switching to choline-free/low-methionine (0.17%) high-fat diet (cdHFD, 9% protein, 60% fat, 2% cholesterol, and 31% carbohydrate, E15673-94, Ssniff, Germany) for two weeks. NASH and control mice (on NC) were sacrificed by cervical dislocation at 6-hour intervals and tissues flash-frozen in N_2_ and stored at −80°C. Animal experiments were ethically approved by the Animal Health and Care Committee of the Government of Schleswig-Holstein and in line with international guidelines for the ethical use of animals. Even numbers of males and females were used for all conditions except for cdHFD at ZT 14 with 1 female and 3 males.

### RNAseq

Total RNA was extracted from livers using TRIzol reagent (Thermofisher, Germany) and columns (Zymo Research, USA) according to the manufactory’s instructions. Trueseq RNAseq comprising 40 million reads per sample (12GB) was performed at Novogene (Germany). For detailed information see supplemental material.

### Differentially expressed gene (DEG) analysis

To identify DEGs independent of sampling time, we used DESeq2 (22) with an adjusted p-value of 0.01 and a log2 fold-change of ±1 as cut-offs pooling samples from all time points (ZTs). For time point specific DEGs, DESeq2 was performed separately for each ZT. Data from three independent mouse NASH RNAseq studies (Lee et al., 2021 (23) - GSE162249; Furuta et al., 2021 (24) - GSE164084; Ægidius et al., 2020 (25) - GSE195483) were downloaded from GEO (https://www.ncbi.nlm.nih.gov/geo) and processed as described above. Of note, genetic background of mice and diet interventions (HFFC - 40 % fat, 20 % fructose, 2% cholesterol – for 24-36 weeks) were largely comparable between the three studies. They differed, however, in light conditions (LD 12:12 for Ægidius et al., 2021, Furuta et al., 2021 and our study; LD 14:10 for Lee et al., 2021).

### Rhythm analyses

To identify diurnal (24-hour) oscillations in gene expression, we combined four independent algorithms: CircaN (26), JTK cycle (27), Metacycle (28), and DryR (29). CircaCompare was used to compare rhythmic parameters between groups (30). For details see supplemental material.

### Gene set enrichment analysis (GSEA)

Enrichment analyses were performed using Gene Ontology (GO) annotations for Biological Processes with DAVID 6.8 software (31) and cut-offs of 3 genes per process and a p-value < 0.05. REVIGO (reduction of 0.5 and default conditions) was used to reduce redundant calls in GSEA (32).

### Random-sampling DEG evaluation

To estimate the effect of time-controlled sampling on the power of DEG detection, we generated 1,000 data sets without time control by randomly picking combinations of 4 samples from the diurnal profile data. DEG evaluation was performed using DESeq2, as described. In addition, a total of 100 temporal profile datasets for NC and cdHFD mice with 4 sampling times and 3 samples per ZT were randomly generated. Previously identified rhythmic genes (using the whole dataset) were kept constant for each generated dataset and used for CircaCompare analysis. Genes with alteration in rhythmic parameters were merged and classified as circadian profiling DEGs (cpDEGs). Comparison between different classes of DEGs was made by using permutation analysis (lmPerm, permutation “Exact”). A p-value of < 0.05 was considered significant.

### Additional experimental procedures

Detailed information for other methods can be found in the supplementary material.

### Data handling and statistical analysis

Samples were only excluded upon technical failure. Analyses were done in Prism 9.4 (GraphPad) with p-value cut-offs of 0.05. 2-group comparisons were done by *Student’s* t-test with Welch correction or Mann-Whitney test for parametric or non-parametric samples, respectively. Normality evaluation was performed using Shapiro-Wilk test. All transcriptome analyses were conducted using R 4.2.1 (R Foundation for Statistical Computing, Austria) or in Prism 9.4 (GraphPad). Used R packages are described in the supplemental material.

## RESULTS

### Time stamping of tissue sampling improves detection of differentially expressed genes in NASH

Our initial step was to evaluate how consistent DEGs were identified between different studies using similar NASH mouse models. Three independent studies using HFFC-induced NASH in C57BL/6J mice were chosen (23–25). RNAseq data were re-analysed with identical parameters in DESEq2 (22). 5,608 NASH-dependent DEGs were identified across all three studies, but only 688 (12 %) of these were identified at the same time in all three (Fig. 1a; Table S1). Considering the marked rhythmicity of the liver transcriptome and liver metabolism (11,33) one explanation for this lack of consistency would be differences in sampling times and control thereof between studies.

**Figure 1:**
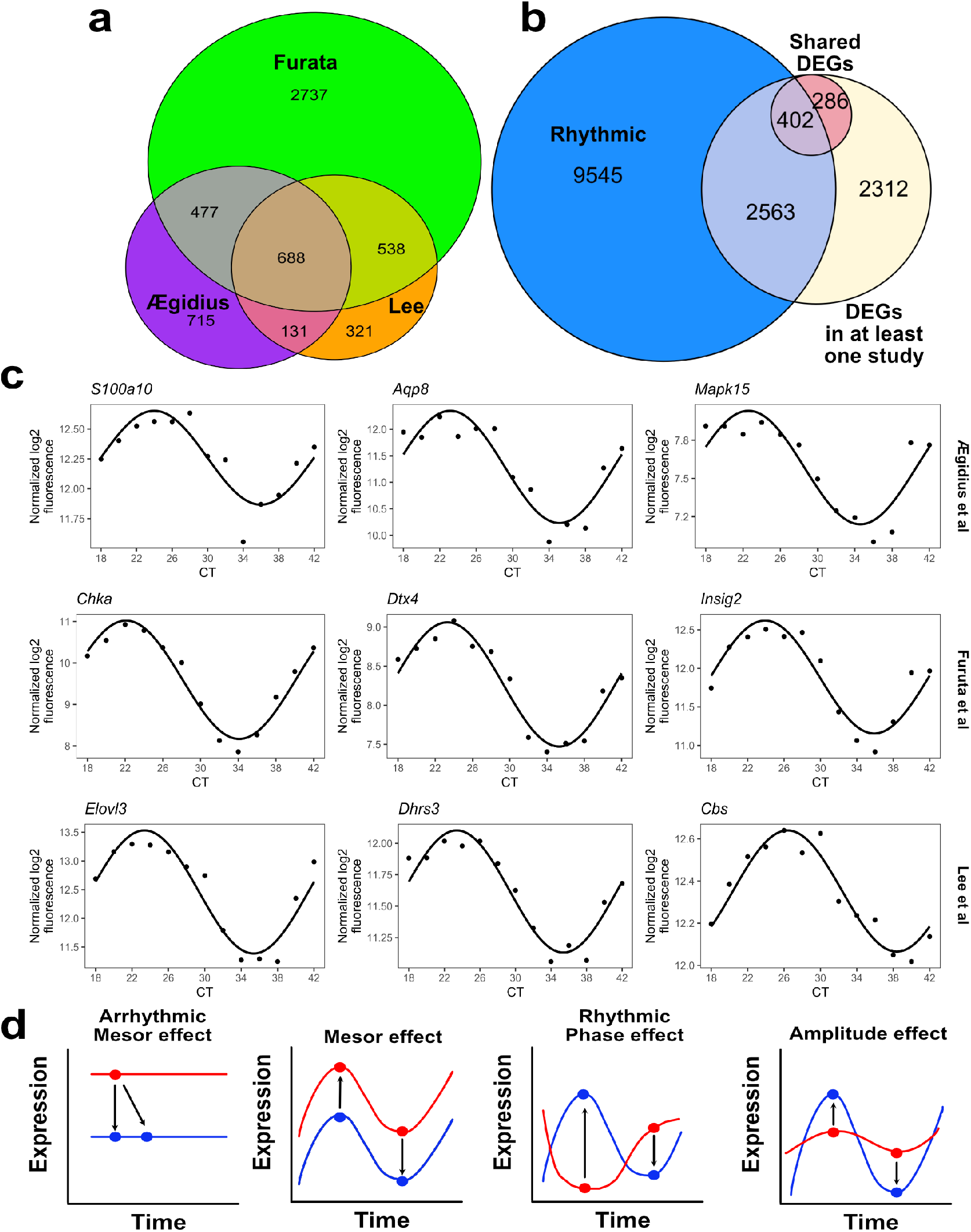
Differentially expressed genes (DEGs) of similar NASH studies show a remarkable small concordance. a) Venn diagram represents the DEGs of three independent studies analyzed using DESeq2 (adj p value < 0.01 and fold change > ± 1). b) Venn diagram shows the number of DEGs (identified in a) that are rhythmic according to liver circadian transcriptome collected every 2 h during 48 h. Shared DEGs (688) were identified in by evaluating on three independent studies. c) representative circadian profiles of exclusive detected DEGs in each dataset. d) blue and red curves represent hypothetical scenarios for changes in mesor, phase, or amplitude, which is described in each plot. Representative scenarios of possible combination in rhythmic parameters that can contribute to the lack of consistent DEGs identified in different studies.

We compared identified DEGs against the Circa database of circadian gene expression (11) and, indeed, 52 % of the 5,608 DEGs detected across studies and 58 % of the shared 688 DEGs were rhythmic (Fig. 1b, c; Table S1). Three principle scenarios are possible in which rhythmicity of gene expression may influence results in a non-time-controlled test scenario (Fig. 1d): if a gene is stably (i.e., without rhythm) expressed under both conditions (Fig. 1d, left panel), then tissue sampling time will have no effect on the outcome of the comparison. However, once DEG expression is rhythmic in any of the two conditions, alterations in mesor (baseline expression), amplitude (variation across time), and acrophase (time of peak expression) may lead to different results depending on the time of sampling (Fig. 1d, right panels). This prompted us to study NASH-induced changes in liver gene expression by profiling transcriptome regulation across 24 hours.

### Effects of NASH on the diurnal regulation of the liver transcriptome

Mice of both sexes were kept on cdHFD for two weeks for rapid NASH induction (34). Liver samples from cdHFD and NC control animals were collected at 6-hour intervals over 24 h and subjected to RNAseq (Fig. 2a). Initially, comparable to previous studies, we evaluated DEGs irrespective of sampling time by pooling data for each condition. This approach yielded 1,684 DEGs (1,214 up- and 470 downregulated; Table S2). When DEGs analysis was done for each timepoint (ZT) separately, 1,664 time dependent DEGs (tdDEGs) were detected, 80 % of which overlapped with the DEGs detected in the previous studies (Fig. 2b; Table S2). Amongst the tdDEGs, 337 genes were consistently up- or downregulated at every single time point in NASH (Fig. 2c). Comparison of these “robust” DEGs with DEGs from previous studies (Fig. 1) revealed a high overlap of 93 % (314 genes). 181 robust DEGs were detected in all datasets (Table S2) constituting a set of bona fide transcriptional NASH markers. Gene set enrichment analysis (GSEA) of robust DEGs revealed several NASH-relevant biological processes such as sterol, lipid, and cholesterol metabolism, inflammation, and insulin signalling (35) (Fig. 2d and examples in e, Table S2).

**Figure 2:**
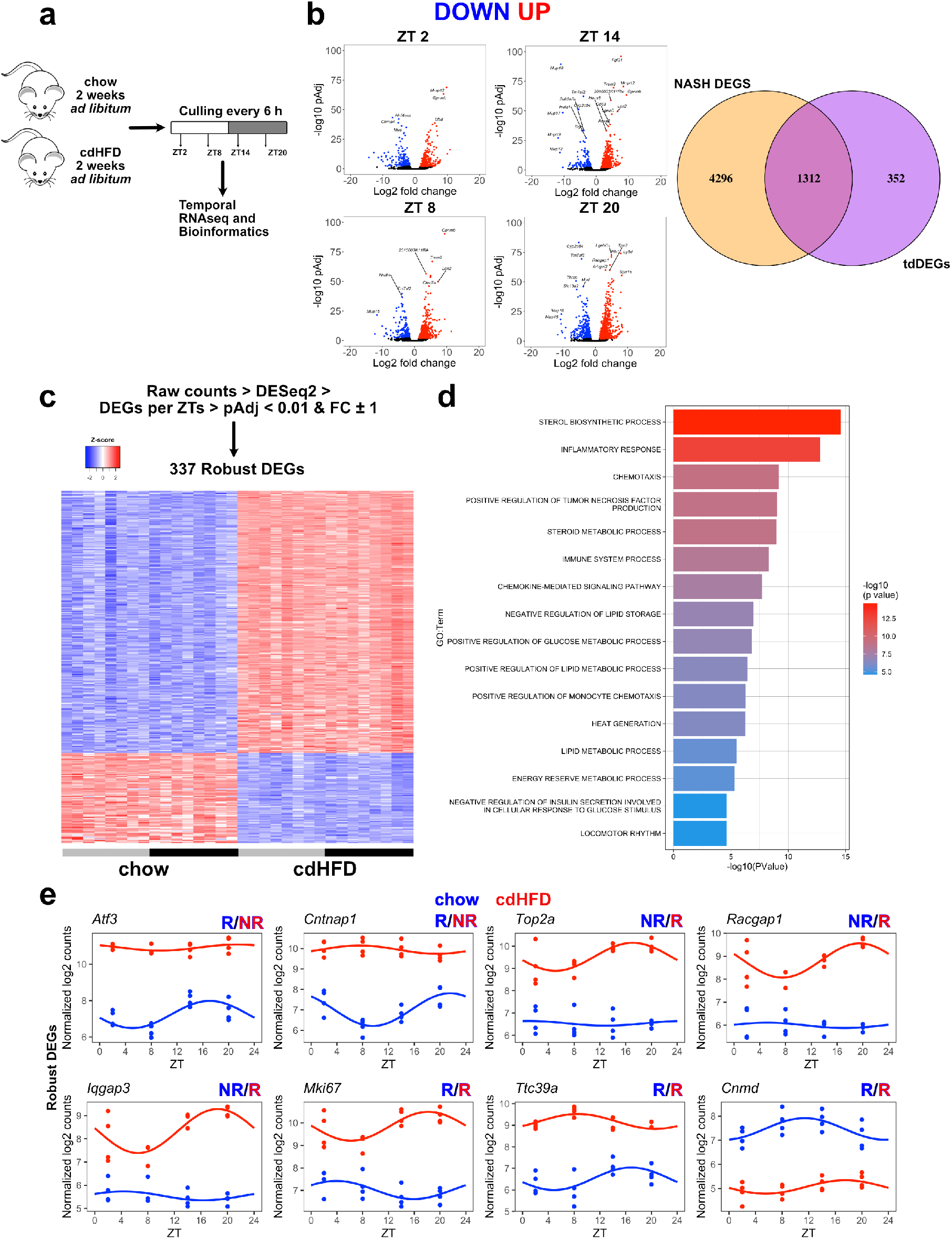
Choline-deficient and methionine low high fat diet (cdHFD) results in marked liver DEGs pattern that is comparable with HFFC diet. a) experimental design is represented. b) DEGs analysis was performed for each ZT and are shown in individual volcano plots. Genes identified in at least one ZT were merged into a single list and classified as time dependent genes (tdDEGs). Venn diagram shows how many genes are exclusively and shared between the NASH DEGs and tdDEGs. c) Robustly DEGs were identified as being differentially expressed across all timepoints (ZTs) and are represented as a heatmap. d) GSEA of robustly DEGs. e) Diurnal profile of robust DEGs that show changes in mesor, amplitude, and/or phase. n = 4 for each ZT. Presence and absence for rhythmicity is shown as R or NR, respectively.

### NASH phase-advances liver transcriptome rhythms

5,151 and 4,215 genes showed significant circadian rhythmicity in either NASH or control livers, respectively, but only 1,714 of these retained rhythmicity across both conditions (Fig. 3a; Table S3). Only slight differences in acrophase distribution between groups were found with the highest number of genes peaking around the night-day transition (ZT23; Fig. 3b). Remarkably, a marked phase advance of about 3h on average was observed in the rhythms of robustly rhythmic genes in NASH (Fig. 3c). This phase advance was mirrored in a marked shift in circadian clock gene expression rhythms (Fig. 3d). To distinguish between NASH and high-fat diet (HFD) effects, we compared cdHFD data to a previously published HFD data set (10). On average, core clock gene rhythms were reduced for mesor, phase and amplitude under cdHFD conditions, and these effects were not observed (mesor, amplitude) or much less expressed under HFD suggesting NASH specific responses (Fig. 3 e – f; Table S3).

**Figure 3:**
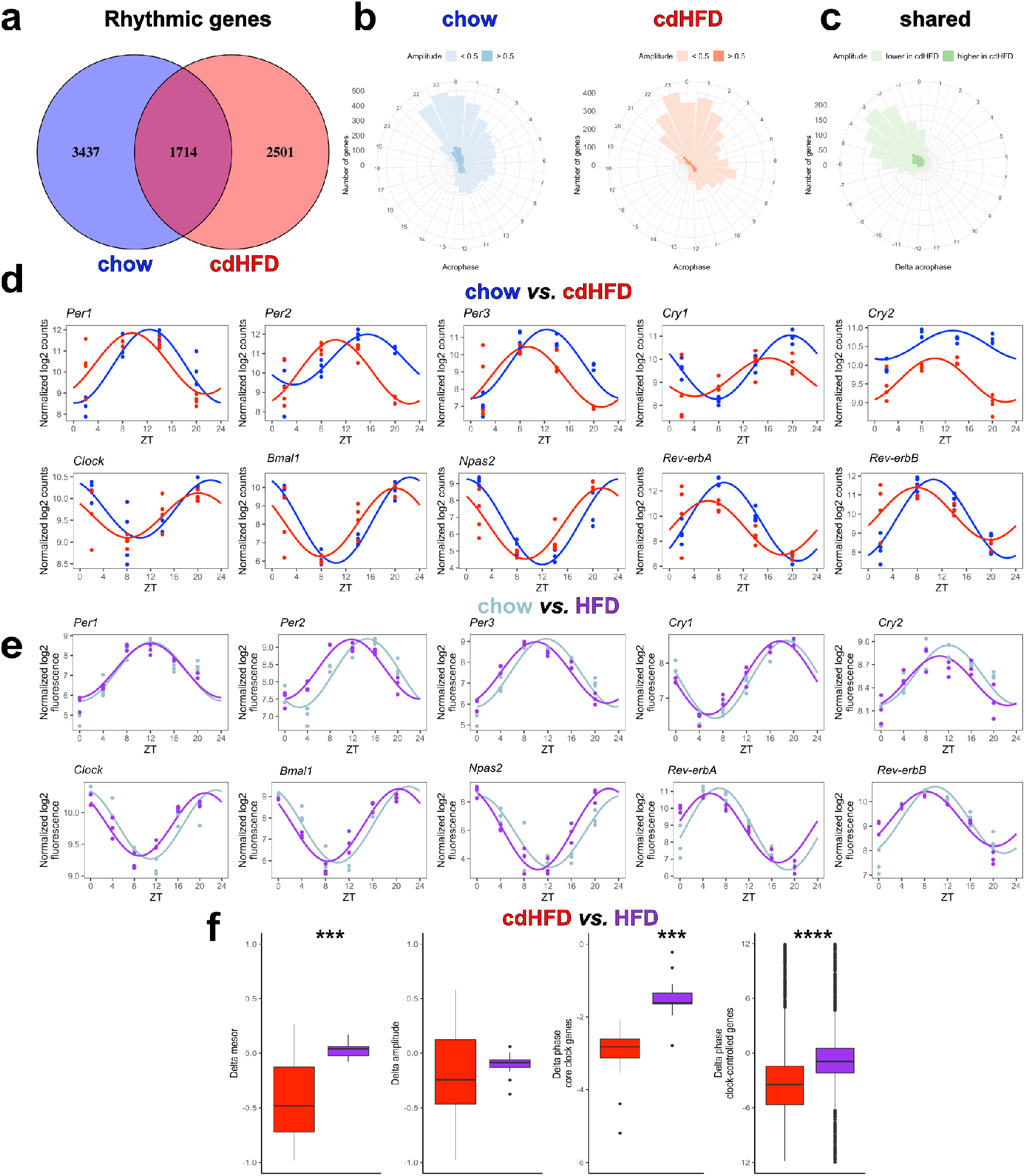
cdHFD results in a marked phase advance in the liver, which is stronger compared to regular HFD. a) Venn diagram shows the number of rhythmic genes for each group. Rhythmicity was performed using CircaN, JTK cycle, Metacycle, and DyrR (p < 0.01). b – c) Roseplots represents the acrophase of chow, cdHFD, and shared rhythmic genes. d – e) Diurnal profile of chow and cdHFD livers and reanalysis of Eckel-Mahan et al 2013 using our bioinformatic pipeline. f) Rhythm parameter evaluation between cdHFD and HFD models. For core clock gene evaluation, paired Mann-Whitney was used. For chow and cdHFD, n = 4 for each ZT. For chow and HFD, n = 3 for each ZT. *** p < 0.001, **** p < 0.0001.

The comparable NASH induced phase advance in the expression profiles of core clock and rhythmic genes suggested a liver clock mediated process. To further evaluate this, we extracted direct targets of core clock transcription factors (TFs; BMAL1, NR1D1, CLOCK, DBP/EBP4) using the ChipSeq atlas database (36). We identified 294, 1,067, 70, and *358 bona-fide* targets of BMAL1, NR1D1, CLOCK, and DBP/EBP4, respectively, which were also rhythmic in at least one diet condition and showed alterations in mesor, amplitude, or phase (Table S4). Over all these clock TF targets, reductions in mesor and amplitude as well as robust phase advances were observed (Fig. 4a). Exclusive targets were filtered and selected for each clock TF and analysed for phase responses. Supporting our assumption, a marked phase advance (of 4 h on average) was observed (Fig. 4 b – d; Table S4). Together these data suggest that NASH induction phase advances the liver clock and, subsequently, clock-controlled genes.

**Figure 4:**
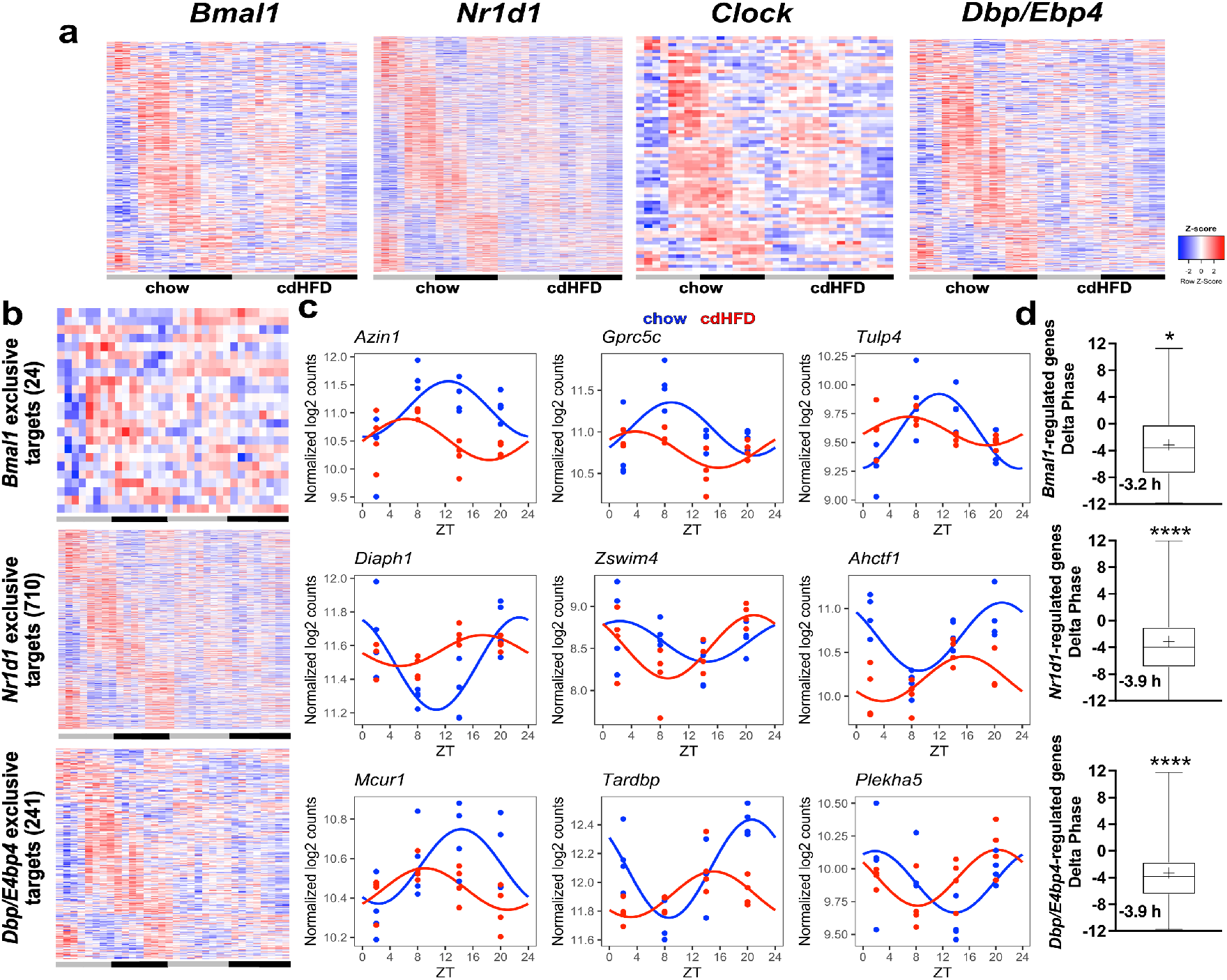
Transcriptional targets of clock transcription factors show a marked phase advance. a) Putative targets of core clock proteins were extracted from Chipseq Atlas (except *Dbp/E4bp4* – see material and methods) and filtered for rhythmicity according to CircaCompare method. Heatmaps of each transcription factor is plotted. b) Exclusive targets for each transcriptional factor were filtered and are shown in heatmaps. c) Representative genes with marked phase advance are depicted. d) Overall phase distribution of the exclusive targets according to CircaCompare. For chow and cdHFD, n = 4 for each ZT.

### NASH induced damping and phase advance of lipid metabolic pathways

To assess the metabolic effects of transcriptome rhythm alterations in NASH, we quantitatively evaluated changes in circadian rhythm parameters in NASH compared to control livers. Most rhythmic genes were altered in mesor (n = 5,062) with similar numbers for increase and decrease. 1,292 genes showed amplitude changes, and most of them (90 %) were dampened in NASH. 1,317 genes showed significant alterations in phase, of which 90 % were phase advanced in line with our previous observations (Fig. 3 c, Fig. 5a – b; Table S5). GSEA of genes with rhythm alterations mainly yielded processes associated with cell damage repair (e.g., GO categories cell cycle progression and DNA repair) and processes associated with lipid metabolism (e.g., lipid and fatty acid metabolism) amongst others (Fig. 5c – Table S5). A marked reduction in mesor and amplitude combined with phase advances was seen for key genes involved in lipid metabolism such as *Lipc*, *Dgat2*, *Acaca*, *Fasn*, *Crot*, *Elovl1/5/6*, *Cpt1a* (Fig. 5d; Table S5). In line with this, NASH led to loss of rhythmicity (amplitude reduction) in liver triglyceride (TAG) levels in addition to a marked 11-fold overall upregulation (Fig. 5d).

**Figure 5:**
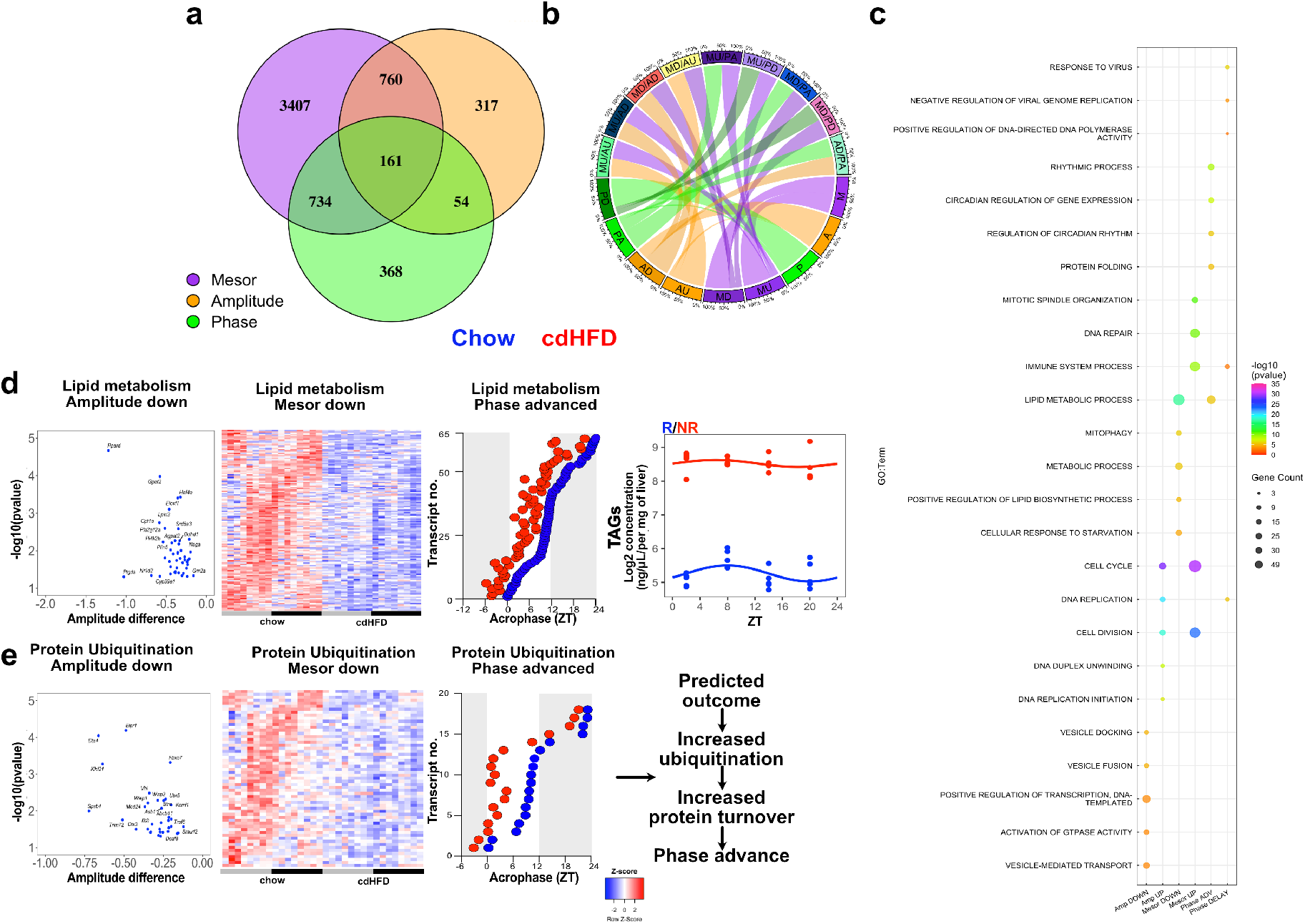
cdHFD leads to a marked diurnal transcriptome rewiring with several metabolic biological processes affected in mesor, amplitude, or phase. a) Venn diagram shows the changes in rhythm parameters detected by CircaCompare method. b) Chord plot represents the combination in rhythmic parameter alterations identified by CircaCompare. c) Top 5 enriched biological processes are represented by each rhythmic parameter. d – e) Genes related to lipid metabolism and protein ubiquitination processes are shown in heatmaps for changes in mesor. Volcano and dot plots show the amplitude and phase change for each gene that participated in a biological process, respectively. Phase was estimated according to CircaCompare. When phase was higher than 18 in the cdHFD group, phase value was subtracted from 24 and plotted to emphasize phase alteration. Liver levels of TAG are shown in log2 values. Presence and absence for rhythmicity is shown as R or NR, respectively. For RNAseq data, n = 4 for each ZT. For TAG data, n = 4 – 5 per ZT.

Interestingly, GSEA also detected protein ubiquitination as a NASH regulated pathway with alterations in mesor, amplitude, and phase. Ubiquitination of circadian clock proteins is a major regulator of clock period and phase (37). In NASH, genes associated with protein ubiquitination displayed reductions in amplitude and/or mesor. For others, marked phase advances were observed (*Ube2h*, *Ube4b*, *Ube2q1*, *Ube2q2*, *Ube2u*) (Fig. 5e; Table S5). These data indicate a time-dependent regulation of protein ubiquitination in the healthy liver, which is altered by NASH. They also suggest a potential direct mechanism for NASH effects on liver clock regulation.

In sum, our comparative rhythm analysis revealed distinct alterations in biological processes associated with liver steatosis. NASH related changes in ubiquitination transcript rhythms may contribute to the phase advance effect observed for clock and clock-controlled gene expression. Further experiments are necessary to validate this premise.

### Temporal control of tissue sampling augments DEG detection

Finally, we compared the power of temporal profiling against traditional one-time point approaches without temporal control. From the previously identified NASH DEGs (n = 5,608 detected by at least one of three studies (Fig. 1), CircaCompare analysis confirmed 1,875 genes with alterations in at least one rhythm parameter (Fig. 6a). Of these, 1,746, 356, and 361 genes showed changes in mesor, amplitude, or phase, respectively. In addition, our profiling approach detected 4,157 DEGs with alterations in one or more rhythm parameters (CircaCompare; n = 3,926) or overall expression (“global DEGs”; n = 231). 568 (83 %) of the DEGs consistently detected by all three previous studies were also detected as altered by this approach. Some examples for NASH induced changes in expression profiles are depicted for alterations in mesor (upper row), amplitude (middle row), and phase (lower row in Fig. 6 b; full list in Tables S5 and S6). Overall, this combined approach outmatched traditional DEG detection for rhythmic parameter changes by a factor of three.

**Figure 6:**
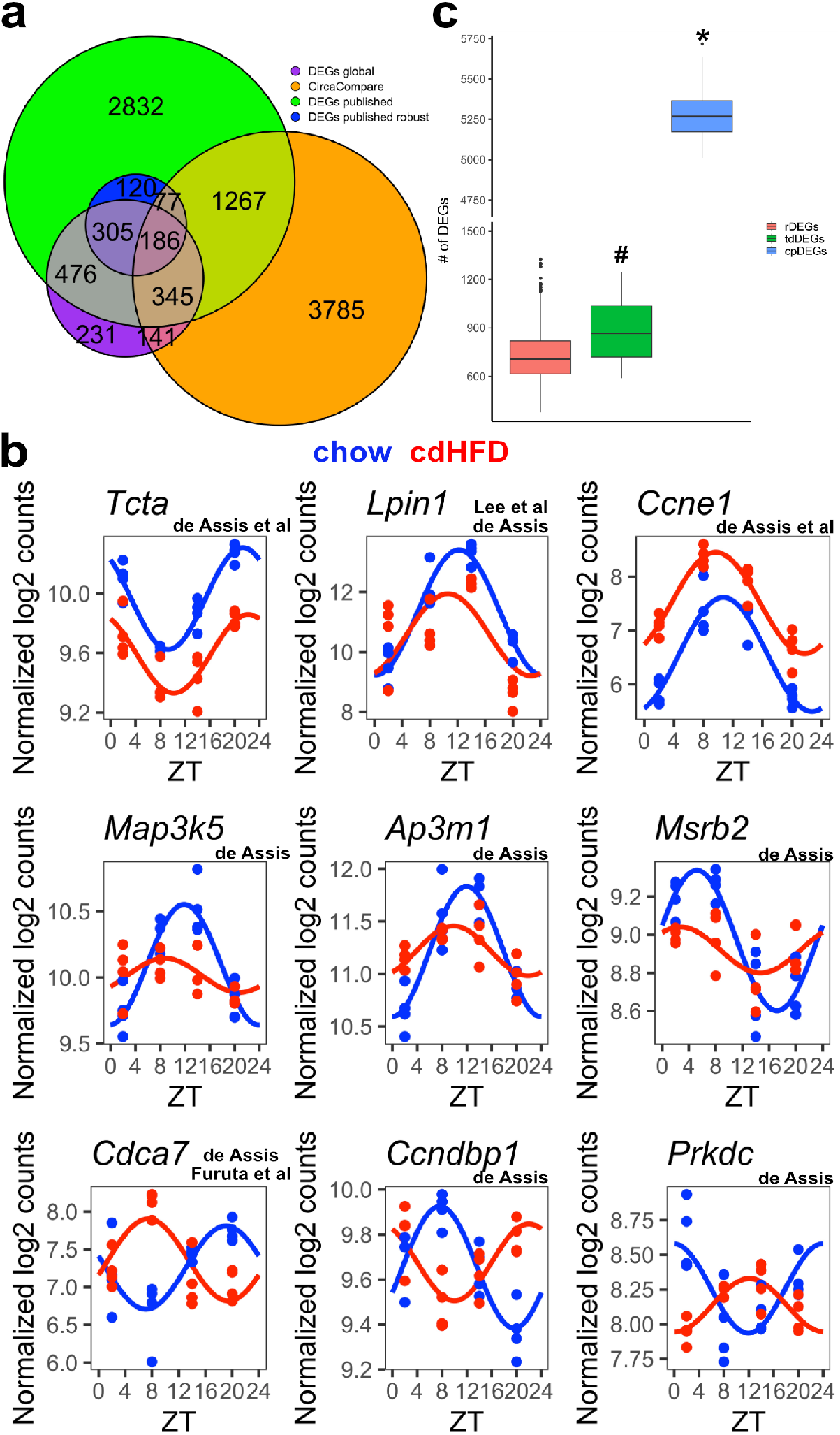
Accounting for sampling time results in increased DEG detection and higher reproducibility of NASH models. a) Venn diagram represents the number of DEGs identified in at least one published study (DEGs published), DEGs identified in all three studies (DEGs published robust), DEGs identified in our dataset disregarding sampling time (DEGs global), and DEGs identified by CircaCompare. b) Representative plots for DEGs that show mesor (upper row), amplitude (middle row), or phase effect (lower row). On top of each gene is shown the study in which such gene was identified. c) Comparison among DEGs generated by random sampling (rDEGs), time-controlled DEGs (tdDEGs), and time-controlled sampling followed by circadian profiling DEGs (cpDEGs). * represents a statistical difference (p-value < 0.001) against rDEGs and tdDEGs; # represents a difference against rDEGs (p-value = 0.029). For chow and cdHFD, n = 4 for each ZT.

To estimate the gain in DEG detection power achieved by controlling for time in transcriptome analyses, we simulated 1 time point random-time sampling (rDEGs) and time-dependent sampling (tdDEGs) from chow and cdHFD data sets and compared these against circadian profiling on a reduced sampling (n = 3 per time point; circadian profiling DEGs - cpDEGs). On average, a higher DEG detection rate of 20 % was obtained when sampling time was controlled (Fig. 6c). Of note, random sampling did not impact robust DEG detection (from cdHFD dataset) as on average 94 % of robust DEGs were identified independent of sampling time. Combined circadian profiling, however, resulted in a 7-fold increase in DEGs, but coming at the cost of a 3-fold increase in sample sizes compared to 1-time point sampling (Fig. 6c; Table S6).

## DISCUSSION

Our findings corroborate previous NASH studies as gene signature suggestive of increased DNA damage and repair (38), disrupted cell cycle progression, tissue repair, and apoptosis (39), and immune system activation (40,41) were identified. Several genes associated with these processes showed increased amplitudes in their circadian expression rhythms, which suggests a stronger level of temporal control. Processes such as lipid and glucose metabolism showed decreased gene expression (mesor DOWN) in NASH livers, thus suggesting an overall activity reduction of energy metabolic pathways. Genes associated with lipid, glucose, and glycogen metabolism also showed reduced amplitude in their daily rhythms or phase advances. A similar phase advance in core clock genes was found suggesting a clock-mediated mechanism. This idea was further supported by phase shifts in bona-fide direct clock target genes.

One could argue that such phase effects are associated with the high steatosis environment induced by cdHFD. Upon evaluating a previous study that used HFD (10), we indeed identified a phase advance in gene expression rhythms, albeit to a much lesser extent (Fig. 3). Therefore, the presence of high lipid environment indeed contributes to the phase advancement, but such effect is further enhanced by pathologic events in the NASH liver. Proper molecular clock functioning is dependent on cycles of protein synthesis and degradation. The first clock mutant (tau) discovered (42) had a shorter locomotor activity period. A subsequent study identified casein kinase 1ε (CK1ε) as responsible for such event due to its capacity to phosphorylate PER. Increased PER degradation as consequence of defective CK1ε phosphorylation resulted in a shorter period (43,44). Protein degradation is primarily mediated via ubiquitination and targeting to proteasomes. Several clock proteins are subject to ubiquitination, which provides an mechanism for period fine-tuning and phase adjustment (37). We suggest that disrupted ubiquitination signalling may result in increased clock protein degradation, which in turn, can phase-advance the core clock (Fig. 3). Interestingly, increased ubiquitination levels have previously been detected in human NASH and were suggested as marker as well as driver of the NASH pathology (45,46).

The observed phase advance in liver glucose metabolism transcripts may have systemic consequences and trigger a compensatory response to normalize serum glucose levels. Similarly, a phase advance in liver lipid metabolism transcripts may shift systemic energy substrate bioavailability to an earlier time. Previous findings showing that HFD-fed mice display increased food consumption during the light (rest) phase (47) corroborate our findings, despite the differences in diet composition. A systemic phase advance in liver transcript rhythms may result in misalignment between geophysical (external) and internal time, affecting metabolic efficiency. In a second potential scenario such phase advance would be restricted to the liver or the liver and a few metabolic organs. Such internal circadian misalignment was shown for HFD conditions in which shifts are mostly restricted to insulin-sensitive organs (48). Similarly, NASH-induced transcriptome changes could contribute to tissue decoupling, further propagating metabolic pathologies and NASH progression (18). Vice versa, addressing such phase shifts by external zeitgebers could slow-down or prevent NASH, and such applications warrant further investigation.

From a medical standpoint, any changes in rhythm parameters are relevant for the optimization of NASH therapies. Alterations in mesor of expression of a relevant gene indicate that the goal of treatment is to modify overall levels of expression or activity, and this goal may be achieved independent of treatment time. This scenario changes once a gene shows an amplitude or a phase effect. Now, the time of treatment may critically determine the therapeutic outcome. NASH induced phase effects may call for a reduction in activity at one time and an increase at another (Fig. 1d). To avoid such complexity, it may be advisable to instead of targeting specific genes/processes to attempt stabilizing the circadian clock itself and in this way indirectly rectify CCG output (49). Considering the changes found in our study, we suggest that clock strengthening either by a pharmacological approach (50) and/or time restricting feeding (51) may benefit NASH treatment. First chrono-modulated pharmacological approaches have been suggested in NASH treatment (52).

Although time is an important regulatory element for health (3), the control for or evaluation of (day-) time as a variable factor is often neglected in many research fields (6). We hypothesized that a lack of control for sampling time augment inconsistencies in results between different NASH studies that used a similar NASH-inducing model. Upon selecting three independent but similar studies (23–25) only a small fraction of DEGs were consistently identified (12%), and many DEGs detected by either one of these studies showed rhythmic regulation of expression across the day (11). To estimate the impact of not having temporal control upon sampling, we determined DEG yields using randomized sampling under time-controlled and -ignorant conditions and compared these against circadian profiling. We confirmed that lack of proper temporal control hampers the detection of DEGs (Fig. 6). Moreover, by performing circadian rhythm analyses, we identified ca. 7-fold more DEGs. Our findings clearly demonstrate the consequences for not accounting for time, which can result in increased false positives and lack of consistency between similar NASH studies.

The marked NASH effects observed on circadian phase of liver transcripts warrant particular attention. Such phase changes, either delays or advances, could mistakenly be interpreted as up- or down-regulation, depending on the time point of assessment. This, in turn, may affect therapy targets. However, with a circadian perspective, more informed decisions on therapeutic strategies could be made. Clearly, considering circadian rhythms in NASH will provide a more comprehensive understanding of the molecular underpinnings of this decrease with high therapeutic potential beyond the development of novel compounds.

**Table 1:** R Packages used in bioinformatic analysis

| Name | Version | Usage |
| --- | --- | --- |
| Eulerr | 6.1.1 | Venn diagram |
| VennDiagram | 1.7.1 | Venn diagram |
| CircaCompare | 0.1.1 | Graph and rhythm comparisons |
| CircaN | 2.0.0 | Rhythm analysis |
| DyrR | 1.0.0 | Rhythm analysis |
| DESeq2 | 1.32.0 | Differentially expressed genes |
| Gplot | 3.1.3 | Heatmap generation |
| Gplot2 | 3.3.5 | Graph generation |
| GraphPad | 9.1 | Graph generation |
| Circlize | 0.4.14 | Chord plot |
| lmPerm | 2.1.1. | Permutation analysis |
| ggbreak | 0.1.0 | Graph generation |

## DATA AVAILABILITY

RNAseq data will be deposited in the Gene Expression Omnibus (GEO) database during the peer review process and will be available after publication.

## CONFLICT OF INTEREST

All authors declare no competing interests that could have an impact on the study.

## ACKNOWLEDGEMENTS AND FUNDING

This work was supported by grants of the German Research Foundation (DFG) to HO 353-10/1, GRK-1957, and CRC/TR 296 LOCOTACT (TP-13).

## AUTHOR CONTRIBUTIONS

LA, MD, HO conceptualization. LA data curation. LA formal analysis and investigation. LA, MD, and HO methodology. HO funding acquisition, project administration, and supervision. LA and HO writing - original draft. All authors: text review & editing.

## SUPPLEMENTAL MATERIAL AND METHODS

### RNAseq

Samples underwent quality control and only those with RNA integrity number (RIN) higher than 6.5 were used. Messenger RNA was purified from total RNA using poly-T oligo-attached magnetic beads. After fragmentation, the first strand cDNA was synthesized using random hexamer primers, followed by the second strand cDNA synthesis using dTTP for non-directional library. The library was checked with Qubit and real-time PCR for quantification and bioanalyzer for size distribution detection. The clustering of the index-coded samples was performed according to the manufacturer’s instructions. After cluster generation, the library preparations were sequenced on an Illumina platform and paired-end reads were generated. Raw data (raw reads) of fastq format were firstly processed through in-house perl scripts. In this step, clean data (clean reads) were obtained by removing reads containing adapter, reads containing ploy-N and low-quality reads from raw data. At the same time, Q20, Q30 and GC content the clean data were calculated.

Reference genome (GRCm38/mm10) and gene model annotation files were downloaded from genome website browser (NCBI/UCSC/ENSEMBL) directly. Paired-end clean reads were mapped to the reference genome using HISAT2 software. HISAT2 uses a large set of small GFM indexes that collectively cover the whole genome. These small indexes (called local indexes), combined with several alignment strategies, enable rapid and accurate alignment of sequencing reads. All the downstream analyses were based on the clean data with high quality. FeatureCounts was used to count the reads numbers mapped to each gene. Genes containing a sum of reads among two groups lower than 100 were excluded from analysis. Remaining genes were log2 transformed in DESeq2 using *vsd* function (blind = FALSE). A total of 18980 transcripts were considered for subsequent analyses.

### Rhythm analyses

CircaN, JTK cycle, and Metacycle algorithms were run in CircaN algorithm using the default parameters. Exact period of 24 h was used. Significance of rhythmicity was set as p < 0.01 for each method. Rhythmic genes were combined into a single list for each group, followed by rhythm parameters comparison using CircaCompare algorithm (1). For mesor and amplitude comparison, CircaCompare was allowed to fit a sine curve irrespective of rhythmicity thresholds. Phase comparison was only performed in robustly rhythmic genes as described (2).

Datasets from Zhang et al., 2014 (3) and Eckel-Mahan et al., 2013 (4) were extracted, log2 (value +1) normalized, and CircaN, JTK cycle, and Metacycle were used to evaluate rhythmicity. DryR method was not included in this analysis due to high temporal resolution and by adding DryR only led to marginal increase (ca. 10%) in novel gene rhythmicity detection in our RNAseq analysis (6h profile). For Zhang et al., 2014, each timepoint consisted of one sample every 2h. For Eckel-Mahan et al., 2013, each timepoint consisted of 3 samples every 4h. For all comparisons in CircaCompare a p value of 0.05 was used. For heatmap and rose plots, phase estimation was extracted from CircaCompare method. Rhythmic gene visualization was made using CircaCompare algorithm.

### Reanalysis of Chipseq analyses

Target genes of transcriptional factors were obtained using the Chipseq Atlas (5). Putative targets genes were selected by selecting genes up to 10K from the transcription start sites (TSS) of mice (mm9 or mm10). Targets genes were filtered for redundancy and only those with an average score > 250 were selected for rhythmicity evaluation using CircaCompare dataset. For DBP/E4bp4 data was extracted from the original study (6). All putative targets were analysed against our rhythmicity analysis dataset.

### Triglyceride (TAG) evaluation

TAG evaluation of tissue was processed according to the manufacturer’s instructions (Sigma-Aldrich, MAK266 for TAG).

## Notes

### Competing Interest Statement

The authors have declared no competing interest.

